# Maternal Obesity Reprograms Differentiation Trajectories of Fetal Hematopoietic Stem and Progenitor Cells Through Altered Inflammatory Signaling

**DOI:** 10.64898/2026.01.29.702548

**Authors:** Brianna M. Doratt, Hami Hemati, Sheridan B. Wagner, Madison B. Blanton, Uriel Avila, Oleg Varlamov, Ilhem Messaoudi

## Abstract

**Background:** Maternal obesity is a global health challenge with profound consequences for offspring health. While its impact on metabolic programming has been widely studied, far less is known about how maternal obesity shapes the fetal immune system. The fetal bone marrow (FBM) is the central site of hematopoietic stem and progenitor cell (HSPC) development, and disruptions in this niche can have lifelong effects on immunity, infection susceptibility, and inflammatory disease risk. In this study, we examined FBM hematopoiesis in a nonhuman primate model of spontaneous maternal obesity.

**Methods:** Using spectral flow cytometry, single-cell RNA sequencing, and functional differentiation assays, we mapped progenitor composition, lineage trajectories, and immune function in offspring exposed to maternal obesity compared with lean controls. These complementary approaches allowed us to capture cellular frequencies and transcriptional programs, while trajectory and signaling analyses provided insight into how progenitor maturation and intercellular communication are disrupted by maternal obesity.

**Results:** Our findings reveal that maternal obesity decreases CD34+ HSPCs and common lymphoid progenitor populations, while expanding megakaryocyte-erythroid and granulocyte-monocyte progenitors. Pseudotime analysis demonstrated altered maturation, with cells accumulating at early differentiation states. Transcriptional profiling uncovered a strong inflammatory bias, with myeloid progenitors upregulating alarmins, interferon-stimulated genes, and proinflammatory mediators. Functionally, monocytes derived from obese FBM showed impaired migratory and colony-stimulating capacity, coupled with exaggerated TNFα responses to LPS stimulation.

**Conclusion:** Together, these results demonstrate that maternal obesity, even in the absence of obesogenic diet, disrupts fetal bone marrow hematopoiesis by altered HSPC maturation, reprogramming lineage trajectories, and inducing inflammatory bias.

## Introduction

The prevalence of obesity (body mass index (BMI) > 30kg/m^2^) among adult women in the Unites States has risen dramatically between 2021 and 2023, with 41.2% classified as obese and 12.1% classified as severely obese (BMI > 40kg/m^2^) (1–3). Maternal pre-pregnancy (pregravid) obesity is associated with a range of obstetric complications, including an increased incidence of gestational diabetes, preeclampsia, cesarean delivery, and spontaneous pregnancy loss (4–8). Several studies have shown that pregravid obesity impacts fetal development and is associated with adverse newborn outcomes, including preterm birth, low standardized assessments of neonatal status (Apgar) score, congenital anomalies, and increased rates of neonatal intensive care unit admission (9–11). These adverse newborn outcomes can persist into early adulthood, with increased risks of allergies and asthma, metabolic disorders, and severe complications from infections (12–14). Animal models of diet-induced maternal pregravid obesity have shown that exposure to pregravid obesity in full-term offspring increases neonatal susceptibility to bacterial and viral pathogens, skews antibody production toward IgE rather than IgG, and drives tissue-specific alterations in immune cell frequencies and pathogen responses (15–17).

Many of the adverse outcomes observed in offspring exposed to pregravid obesity can be linked to altered immune system maturation and function (18, 19). Clinical studies have shown that pregravid obesity alters immune cell frequencies in umbilical cord blood (UCB) and impairs their responsiveness to stimulation (20–23). Mechanisms governing these changes have been tied to epigenetics and transcriptional alteration within monocytes and CD4+ T cells, but our understanding remains incomplete (20, 22). In the late-term fetus and postnatal offspring, nearly all immune cells originate from hematopoietic stem and progenitor cells (HSPCs) residing in the fetal bone marrow (FBM), with the exception of tissue resident immune cells that are seeded earlier in gestation from the fetal yolk sac and fetal liver (24, 25). Hematopoiesis, the differentiation of immune and other blood lineages from HSPCs, is a dynamic process supported by the FBM microenvironment (26). Previous reports have demonstrated that pregravid obesity can impact HSPCs and the FBM niche. For example, murine models of high-fat diet (HFD)-induced pregravid obesity have shown reduction in HSPCs in the fetal liver, expansion of terminally differentiated myeloid and B cells, and transcriptional alterations in pathways including metabolism, glucose tolerance, inflammation, and signaling (27–29). Furthermore, in the NHP model of maternal Western style diet (WSD)-induced obesity, we previously demonstrated premature development of bone marrow adipocytes, reduced abundance of lymphoid progenitors, and a proinflammatory transcriptional landscape in FBM primed for innate immune responses (30, 31).

Although significant progress has been made in understanding how pregravid obesity affects fetal and neonatal immunity, the specific mechanisms driving altered hematopoiesis within the FBM niche remain poorly defined. Furthermore, murine and NHP models of maternal obesity to date have primarily relied on high-fat, calorically rich WSDs to induce the condition (32, 33). However, obesity can also result from genetic predisposition, which affects appetite and basal metabolism, as well as from physical inactivity, environmental influences, or comorbid medical conditions (34, 35). Therefore, we extended our previous studies and examined alterations in HSPCs and their differentiation within the FBM of gestational day (GD) 130 fetuses derived from lean or obese rhesus macaques fed a regular chow diet. Rhesus macaque gestation and fetal development closely mirrors human development (36, 37), with GD130 representing the mid-third trimester, which typically spans 110-165 days (36, 37). We report that diet-independent maternal obesity alters the fetal bone marrow environment by inducing a proinflammatory transcriptional landscape, promotes myelopoiesis, and impairs the ability of differentiated immune cells to respond to antigen.

## Methods

### Cohort description

Female rhesus macaques underwent time-mated breeding at the Oregon National Primate Research Center (ONPRC). All animals in this study were managed according to the ONPRC animal care program, the AAALAC International, and guidelines set forth by the United States Department of Agriculture. Animals received ad libitum access to fresh water and food (Purina 5000 Fiber-balanced Monkey Diet, Purina Mills, Richmond, IN, USA) supplemented with fruits and vegetables, as part of the environmental enrichment program established by the Behavioral Services Unit. The nutritional plan utilized by the ONPRC is based on National Research Council recommendations. In the present study, 6 lean and 7 spontaneously obese 8-to 12-year-old female rhesus macaques maintained on a regular monkey chow diet were selected from the ONPRC colony based on the body condition score (BCS). BCS is a semiquantitative method of assessing body fat and muscle, on a scale of 1-5, by palpation of key anatomic features including the hip bones, facial bones, spinous processes and ribs as well as muscle mass present over the hip bones and prominence of the ischial callosities (38). Obese animals were defined as having a BCS ≥4, with lean animal having a BCS of 2-3.

### Sample collection and processing

Fetuses were obtained via scheduled cesarian section between GD130 and GD135 and FBM mononuclear cells were isolated as described previously (30). Briefly, both fetal femurs were cleaned by removing soft tissues and cartilage and manually crushed with a ceramic pestle in 10 mL of cold RPMI supplemented with 3% BSA, 1% Pen/Strep, 1% L-glutamine, and 10mM HEPES. Cell suspension was filtered through a cell strainer and centrifuged over a Ficoll gradient. FBM mononuclear cells residing at the Ficoll interface were carefully collected and red blood cells were lysed. Cells were washed twice with media, cryopreserved using CryoStor10 (StemCell Technologies) and stored in a liquid nitrogen cryogenic unit for long-term storage.

### Spectral flow cytometry

Spectral flow cytometry was performed as described previously (39). Briefly, cells were stained (n= 6 lean, 7 obese) with a surface antibody cocktail in the presence of 50 µl Brilliant Stain Buffer, 5 µl TruStain FcX, and 5 µl True-Stain Monocyte Blocker for 30 minutes at 4°C in the dark. Following surface staining, cells were washed and permeabilized using the Tonbo permeabilization buffer (Tonbo Biosciences) for 2 hours at 4°C and incubated with intracellular antibodies overnight at 4°C in the dark. Samples were analyzed using a Cytek Aurora spectral flow cytometer equipped with five lasers and SpectroFlo Software (v3.0). Spectral unmixing was performed using single-stained reference controls (beads or cells), with autofluorescence extraction enabled. Unmixed FCS files were analyzed using FlowJo (v10.10.0). Initial quality control and removal of anomalous events were conducted with FlowAI (v2.3.2) (40).

The full panel of antibodies consisted of: CD3 (BUV496, OKT3, BD Biosciences), CD20 (BUV496, 2h7, BD Biosciences), CD14 (BUV805, MSE2, BD Biosciences), CD45RA (BUV395, 5H9, BD Biosciences), CX3CR1 (RB780, 2A9-1, BD Biosciences), NF-κB p65 (pS529; PE, Kl 0-895.12.50, BD Biosciences), CD184 (CXCR4; BV480, 12G5, BD Biosciences), HLA-DR (Spark Violet™ 538, L243, BioLegend), CD16 (APC/Fire™ 750, 3G8, BioLegend), CD34 (APC/Cyanine7, 561, BioLegend), CD282 (TLR2; PE/Cyanine7, TL2.1, BioLegend), CD115 (CSF-1R; PE/Dazzle™ 594, 9-4d21ef, BioLegend), SPI1 (PU.1; Alexa Fluor 647, 7C6B05, BioLegend), CD64 (VioBlue, 10.1.1, Miltenyi Biotec), TNF RI/TNFRSF1A (PE/Cy5.5, H398, Novusbio), CCR2 (APC, 48607, Novusbio), HIF-1 alpha (Alexa Fluor® 488, H1alpha67, Novusbio), CD38 (PerCP or PE/Cyanine7, AT1, Novusbio), CXCR1/IL-8RA (Alexa Fluor® 700, 42709, Novusbio), CD123 (Brilliant Ultra Violet™ 737, 6H6, eBioscience), CD90 (Thy-1; Brilliant Ultra Violet™ 563, eBio5E10 (5E10), eBioscience), CD127 (IL-7RA; Super Bright™ 780, eBioRDR5, eBioscience), CD284 (TLR4) (Super Bright™ 600, HTA125, eBioscience), TNF alpha (Brilliant Violet™ 650, MAb11, eBioscience), IL1R2 (PE, 34141, Thermofisher Scientific), and IRF8 (PerCP-eFluor™ 710, V3GYWCH, eBioscience).

### Liquid Differentiation Assay

The liquid differentiation assay was performed as described previously (41). Briefly, FBM cells was thawed (n=5 lean, 4 obese) and stained with anti-CD34 antibody (PE-Cy7, 563, BioLegend) before sorting on a Sony SH800 Cell Sorter. A total of 1000 sorted CD34+ cells were plated per well of a 96-well plate in 100 μL StemSpan SFEM Media (StemCell Technologies) supplemented with StemSpan Myeloid Expansion Supplement II and incubated at 37°C in a 5% CO_2_ environment. On day 14, cells were harvested for downstream scRNA-seq analysis, phenotyping, and stimulation. Differentiated FBM were phenotyped by surface staining with CD34 (PE-cy7, 563, BioLegend), CD14 (AF700, MSE2, BioLegend), HLA-DR (APC-cy7, L243, BioLegend), CD16 (PB, 3G8, BioLegend), and CD115 (PE, 9-4d21ef, BioLegend). Stimulation of differentiated FBM was performed by incubating cells in the presence or absence of LPS (0.5 mg/mL) for 16 hours. Stimulated cells with the addition of Brefeldin A (BFA, BioLegend) were used for ICS by staining with CD14 (AF700, MSE2, BioLegend) and HLA-DR (APC-cy7, L243, BioLegend). Cells were fixed and permeabilized using the BioLegend Fix Perm kit before intracellular staining with anti-TNFα antibody (APC, Invitrogen). Phenotyping and ICS samples were acquired with an Attune NxT Flow Cytometer (ThermoFisher Scientific) and analyzed using FlowJo software (v10.10.0). Additionally, supernatants from the stimulations incubated without BFA were run on a custom NHP Millipore Luminex kit with analytes BDNF, CXCL13, CCL11/Eotaxin, CCL2, CD40L, CCL20, CXCL10, CXCL11, FGFbasic, G-CSF, GM-CSF, IFN-b, GZMB, IFN-a, IFN-g, IL-1b, IL-10, IL-12, IL-13, PD-L1, IL-6, IL-8, PDGF-AA, PDGF-BB, CXCL2, CCL5, TNF-a, and VEGF.

### scRNA-seq library generation

FBM cells were thawed (n=3 per group), washed in DPBS with 2% FBS and incubated with Rhesus Fc Block. Each sample was stained with anti-CD34 antibody (PE-Cy7, 563, BioLegend) and labeled with a distinct TotalSeq Hashtag Oligo antibody (HTO, BioLegend) per manufacturer’s instruction. Pellets were washed twice in DPBS with 2% FBS and pooled by maternal condition. Propidium iodide was added immediately before sorting and live CD34+ FBM cells were sorted using a Sony SH800 Cell Sorter. Total differentiated FBM (n=3 per group) were labeled with a distinct TotalSeq Hashtag Oligo Antibody (HTO, BioLegend) and incubated per manufacturer’s instruction. Pellets were washed twice in DPBS with 2% FBS and pooled together. Sorted FBM cells or total differentiated FBM were counted in triplicate and suspended in DPBS with 2% FBS to a final concentration of 1600 cells/μL. Cell suspensions were immediately loaded on the 10x Genomics Chromium X with a target of 30,000 cells. Libraries were prepared using V3.1 chemistry for gene expression and 3L Feature Barcode Library Kit per manufacturer’s instructions (10x Genomics). Libraries were sequenced on Illumina NovaSeqX with a sequencing target of 30,000 gene expression reads and 5,000 feature barcoding reads per cell.

### Single-cell RNA-seq data analysis

Raw reads were aligned and HTOs were demultiplexed using Cell Ranger (v7.2, 10x Genomics) against the Macaca mulatta reference genome (mmul10) using the multi-option. Droplets with ambient RNA or potential doublets (<400 or >4000 detected genes) and dying cells (>20% total mitochondrial gene expression) were excluded during initial QC. All libraries were integrated using Harmony before data normalization and variance stabilization were performed using the NormalizeData and ScaleData functions in Seurat (v5). Dimensionality reduction was performed using RunPCA function to obtain the first 30 principal components and clusters visualized using Seurat’s RunUMAP function. Cell types were assigned to individual clusters using FindAllMarkers function with a log2 fold change cutoff>0.4, FDR<0.05, and using a known catalog of well-characterized scRNA markers for NHP leukocytes. Differential Trajectory, Topology, and Progression analysis was performed using Slingshot (42) with the Condiments package (v1.17) (43), with the indicated HSC population rooted as the start. Temporally expressed genes were identified by ranking all genes by their variance across pseudotime and then further fit using the generalized additive model with pseudotime and maternal phenotype as independent variables. The progressionTest function was used with the Classifier method to test if lineages were statistically different by condition. The conditionTest function was used with l2fc = 0.585 for each lineage independently to identify gene-level alterations with condition.

The R CellChat(44) package was employed to infer probable intercellular communication networks, utilizing the CellChatDB.human database. Data was preprocessed with the function identifyOverExpressedGenes and identifyOverExpressedInteractions and communication probabilities between clusters were determined with computeCommunProb (truncatedMean, trim=0.1, interaction.range=250, contact.range=100). Communications were filtered (filterCommunication) to a minimum number of 10 cell and signaling pathway probabilities were calculated (computeCommunProbPathway).

### Statistical Analysis

GraphPad Prism 10 was used for data analysis and error bars indicate the standard error of the mean. Normality testing was performed using the Shapiro-Wilk test and outliers were assessed using the ROUT test with Q=1%. Two group comparisons of normally distributed populations were compared using a Welch’s t-test, while populations that fail normality were compared using the non-parametric Mann-Whitney t-test. Four way comparisons of stimulated and non-stimulated conditions were assessed using a one-way ANOVA with Bonferroni correction. For all figure panels p-values were indicated using the following symbols: # = p-value <0.1, *= p-value <0.05, ** = p-value <0.01, *** = p-value <0.001, **** = p-value <0.0001.

## Results

### Maternal Obesity Alters FBM Composition and Leads to Progenitor Imbalance

To assess how maternal obesity alters the phenotype of FBM HSPCs, we used an 11-color spectral flow cytometry panel to investigate alterations in frequencies of HSCs, various progenitor populations as well as differentiated immune populations (Supp.Fig.1A-B). We observed an increase in frequencies of lineage negative (Lin−, CD3-CD20-CD14-) non-hematopoietic cells in the obese group compared to lean controls (Fig.1A). In contrast, the frequencies of HSPCs, including late CD34^+^CD38^+^ (P1) and early C34^+^CD38^-^ (P2) progenitors, were reduced in FBM from the obese group (Fig.1A). We also observed a significant decrease in hematopoietic stem cells (HSC), while megakaryocyte and erythrocyte progenitors (MEP) were elevated in the obese group relative to lean controls (Fig.1B).

**Figure 1:**
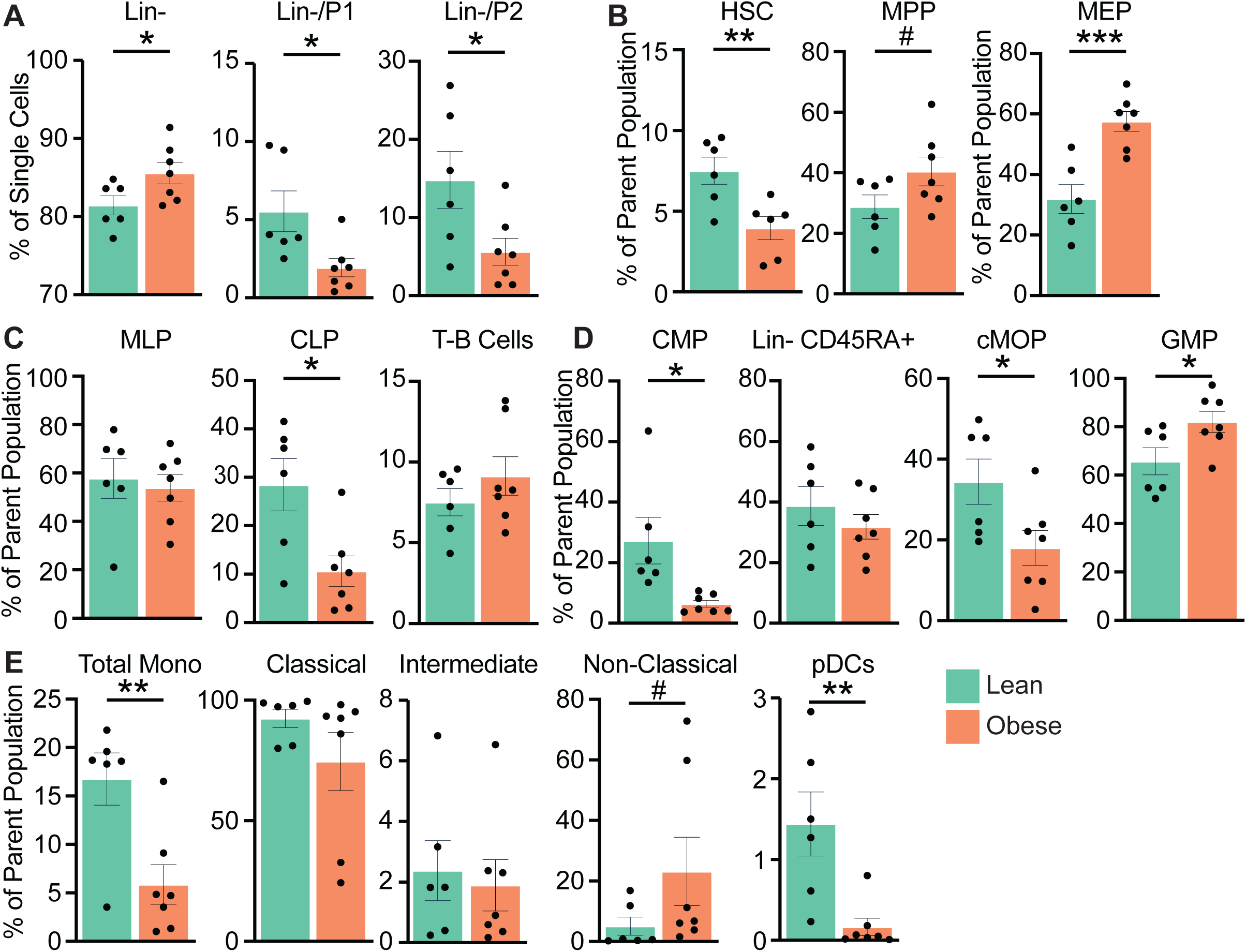
Alterations in frequencies of fetal HSPC and immune cell populations in response to maternal obesity. A-E) Bar graphs showing the indicated progenitor populations, derived either from the single-cell gate or the parent population. Gating strategy shown in Supp Figure 1. A) Lin⁻ (lineage-negative, CD3⁻CD20⁻CD14⁻), P1 (late progenitor, CD34⁺CD38⁺), and P2 (early progenitor, CD34⁺CD38⁻). B) HSC (hematopoietic stem cell), MPP (multipotent progenitor), and MEP (megakaryocyte–erythroid progenitor). C) MLP (multi-lymphoid progenitor), CLP (common lymphoid progenitor), and T cell (T lymphocyte), B cell (B lymphocyte). D) CMP (common myeloid progenitor), cMoP (common monocyte progenitor) and GMP (granulocyte–monocyte progenitor). E) Monocytes (total population, classical, intermediate, non-classical), and pDC (plasmacytoid dendritic cell).

Analysis of the lymphoid lineage showed no difference in the abundance of early multi lymphoid progenitors (MLP) or the terminally differentiated T and B cells (Fig.1C). However, common lymphoid progenitors (CLP) were significantly reduced in the obese group (Fig.1C). Within the myeloid lineage, we observed a reduced frequency of common myeloid progenitors (CMP) with no change in the abundance of Lin^-^ CD45RA^+^ progenitors (Fig.1D). When the Lin⁻ CD45RA⁺ compartment was further subdivided into granulocyte–monocyte progenitors (GMP) and common monocyte progenitors (cMoP), maternal obesity was associated with a shift toward increased GMP and reduced cMoP, suggesting a skewed differentiation trajectory away from more classical monocyte derivation and toward a more “neutrophil-like” monocyte population(45) (Fig.1D). In line with this finding, the analysis of terminally differentiated myeloid cells showed a reduction of monocyte frequency with no change in the relative proportions of classical or intermediate monocytes, and a trend toward increased non-classical monocytes (Fig.1E). In addition, CD123+ plasmacytoid dendritic cells (pDC) decreased in the obese group compared to lean (Fig.1E). Overall, our data demonstrate that maternal obesity disrupts fetal bone marrow hematopoiesis by altering progenitor biogenesis by impairing lineage commitment and/or by attenuating their terminal differentiation.

### Maternal Obesity Shifts HSC Trajectories Toward Early Pseudotime States

To investigate the mechanisms underlying altered HSPC composition and differentiation in the FBM niche associated with maternal obesity, we performed scRNA-seq on fetal CD34+ HSPCs. We identified 10 transcriptionally distinct cell clusters that were consistently represented across both lean and obese groups (Fig.2A). Cell clusters were identified and annotated using established marker genes for HSPCs(30, 46) (SuppTable1). Among these clusters, we identified two HSC populations, with the first population expressing higher levels of genes (*CD38, CD48,* and *CDK6)* associated with HSC activation, early differentiation, and lineage priming, suggesting a more transcriptionally active, differentiation-primed short-term hematopoietic stem cell (ST-HSC) state(46–49) (Fig.2A,B). The second HSC population exhibiting high expression of *FGD5*, *MECOM*, *MEIS1*, and *HLF* transcription factors characteristic of a more quiescent, primitive long-term hematopoietic stem cell (LT-HSC) state(46–52) (Fig.2A,B). Two multipotent progenitors (MPP) clusters were also identified, both marked by expression of *ELANE* and *CLEC12A* and lower *CD34* levels. One of these clusters, Prolif_MPP, showed high proliferative activity, as indicated by higher *MKI67* expression (Fig.2A,B). Other progenitor clusters included monocyte dendritic cell progenitors (MDP; *AZU1*, *MPO*, and *LYZ*), myeloid progenitors (MyeloP; *LYZ*, *S100A9*, *FCGR1A*, and *MYD88*), MEP (*VWF*, *TPM1*, and *GATA1*), and erythroid progenitors (EryP; *KLF1, GPX4, TFRC, HBA*) (Fig.2A,B). Finally, granulocyte progenitors (GranulP) were identified by *CPA3* expression, while common lymphoid progenitors (CLP) expressed *CD79B*, *DNTT*, *IL7R* and *BCL11A* (Fig.2A,B).

**Figure 2:**
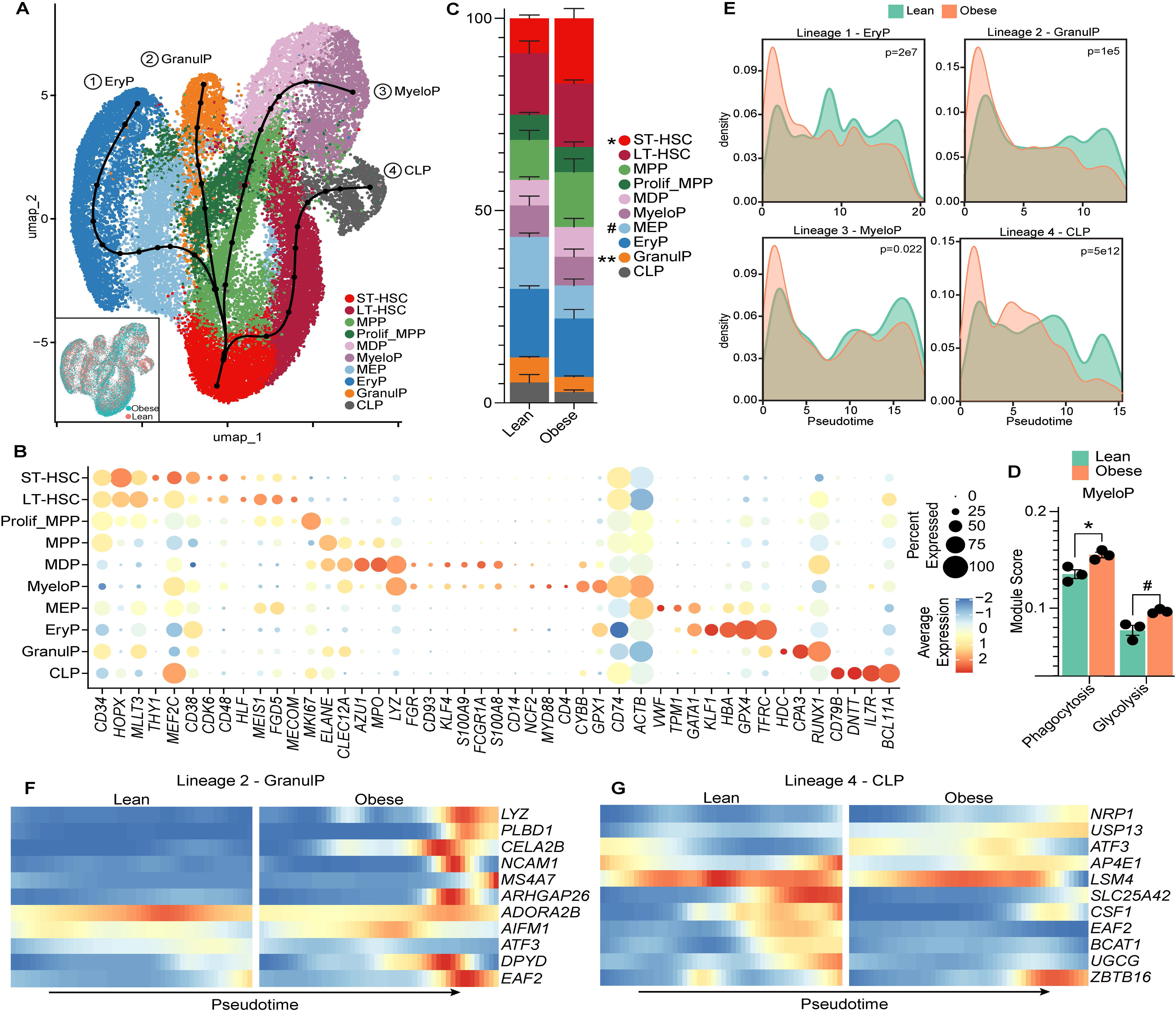
Maternal obesity alters myelopoiesis assessed by scRNA-seq. A) UMAP of 30,785 sorted CD34+ FBM cells (14,238 lean and 16,547 obese) derived from lean (n=3) or obese (n=3) dams. Colors indicate the 10 distinct clusters. UMAP is overlaid with four lineage trajectories. Inlayed UMAP colored by cell origin of fetuses from lean or obese dams, showing proper dataset integration. B) Bubble plot of marker genes used to identify the 10 clusters. Bubble size represents the percentage of cells in each cluster expressing the indicated marker, while bubble color reflects the average expression level of the marker within the cluster. C) Stacked bar plot showing the relative cluster frequencies. D) Bar plot showing the indicated module score for the MyeloP cluster. E) Progression plot depicting cell densities along pseudotime for the indicated trajectories. F-G) Heatmap of the indicated differentially expressed genes from F) Lineage 2 - GranulP (F) and Lineage 4 - CLP1 (G) along pseudotime split by lean and obese.

We next performed a trajectory analysis to assess HSC differentiation. Using the ST-HSC cluster as the root, we identified four distinct differentiation trajectories (Fig.2A). Lineages 1-3 transitioned through MPP and the Prolif_MPP clusters before ending in EryP (lineage 1), GranulP (lineage 2), or MyeloP (lineage 3) (Fig.2A, Supp.Fig2A-C). In contrast, lineage 4 terminated in the CLP cluster (Fig.2A and Supp.Fig.2D). Comparison of cluster frequencies showed a higher proportion of ST-HSC in the obese group, accompanied by decreased GranulP frequency (Fig.2C). Additionally, module score analysis revealed that the MyeloP cluster had a significant increase in phagocytic potential, with a trend toward elevated glycolytic activity in the obese group (Fig.2D). No differences in hypoxic signatures, glycolysis or oxidative phosphorylation (OxPhos) transcriptional signatures were observed in any other cluster. To assess group-specific differences, we compared cell densities along the pseudotime within each lineage (Fig.2E). All four trajectories were significantly altered with maternal obesity, marked by an increased cell density at early pseudotime suggesting altered maturation (Fig.2E), in line with flow cytometry data showing reduced frequencies of mature myeloid cells with maternal obesity (Fig.1E).

### Maternal Obesity Alters HSPC Maturation and Induces Inflammatory Bias Across Lineages

We next examined pseudotime-dependent gene expression changes to uncover mechanisms driving altered differentiation trajectories with maternal obesity. We identified 10 differently expressed genes (DEGs) along the GranulP lineage (SuppTable2). Compared to lean controls, GranulP lineage cells exposed to maternal obesity exhibited increased expression of *AIFM1*, a gene involved in apoptosis and mitochondrial bioenergetics(53), at mid-pseudotime. Expression of *ADORA2B,* an adenosine receptor whose expression in granulocytes has been linked to limiting inflammation by suppressing TNF-α release and reducing oxidative activity(54, 55), decreased across pseudotime in GranulP cells from the obese group (Fig.2F). In contrast, several other genes were upregulated at only the terminal end of the GranulP lineage in the obese group relative to lean group (Fig.2F). These genes include *LYZ,* an antimicrobial enzyme that breaks down peptidoglycan in bacterial cell walls and has been associated with obesity-induced low grade inflammation(56), and *PLBD1*, a lysosomal enzyme-like protein and transcription factor that have a role in degradation processes that effect phagocytosis and antigen processing(57) (Fig.2F).

A total of 36 DEG were identified along the CLP lineage (SuppTable2). Expression of *AP4E1*, an adaptor protein critical for intracellular signaling(58), showed an inverted pattern, with lean group showing high expression at early and late pseudotime stages, while the obese group displayed peak expression at intermediate pseudotime, indicating a reprogramming of lymphoid differentiation kinetics. Additionally, at the terminal end of the lineage, maternal obesity was associated with decreased expression of several metabolic related genes included *SLC25A42*, a mitochondrial coenzyme A transporter that is critical for mitochondrial metabolism(59), and *LSM4*, a component of the spliceosome complex that aids in RNA metabolism and has been correlated with immune infiltrate levels in cancer and infection(60) (Fig.2G). We also observed a decrease in genes associated with lymphocyte development and response including *CSF1*(61), *EAF2*, *BCAT1*, and *UGCG* (Fig.2.G). In the CLP lineage pregravid obesity was associated with upregulated expression of NKT cell markers *NRP1 and ZBTB16* (Fig.2.G). Finally, we observe an increase in the expression of stress inducible transcription factor *ATF3*, as well as *USP13*, a deubiquitinase that regulates cell cycle progression(62, 63) (Fig.2G).

In the EryP lineage, we observed 421 DEGs between lean and obese groups (SuppTable2). Functional enrichment of these DEGs mapped to gene ontology (GO) terms associated with cell activation, inflammatory response, megakaryocyte differentiation and heme metabolism (Fig.3A). At the terminal end of pseudotime, *MAP4K4* and *ISG20* (a marker of interferon activation) expressions levels were markedly increased in the obese group (Fig.3B). In contrast, lineage-specific genes such as *HBQ1*, *HBA*, and *HBE1* were decreased in the obese group (Fig.3B). Finally, we observed reduced expression of the adhesion molecule *PECAM1* and the mitochondrial gene *ATP5IF1* with maternal obesity, accompanied by increased expression of chromatin remodeling genes, including *H2AC20*, *HIST1H2AE*, and *H1-5* (Fig.3B).

**Figure 3:**
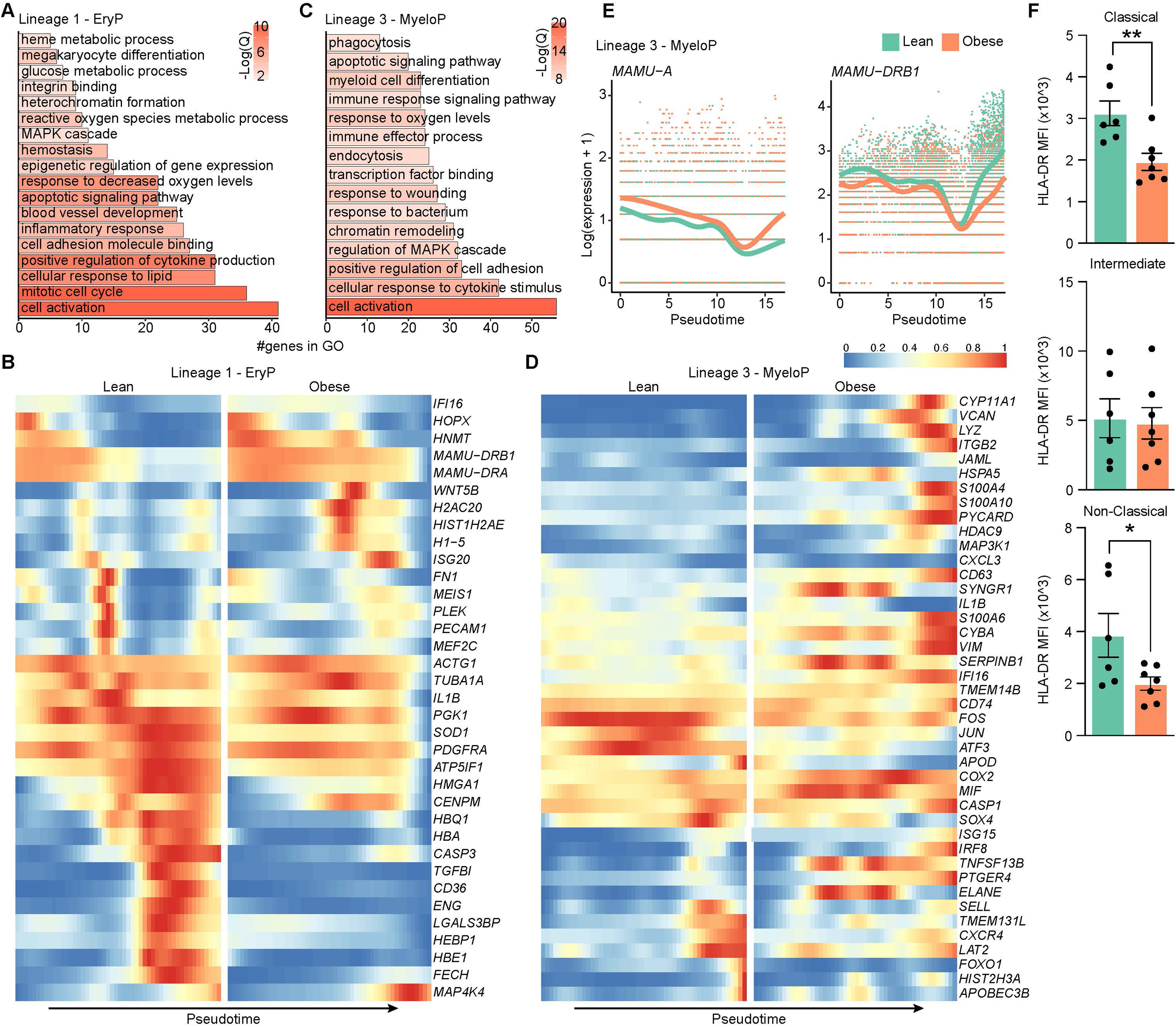
Maternal obesity is associated with higher and earlier expression of inflammatory genes. A, C) Bar plot of gene ontology terms for genes that are differentially expressed along pseudotime in Lineage - 1 EryP (A) and Lineage - 3 MyeloP (C). Bar length represents the number of genes mapping to each term and bar color indicates the -log10(Q-value). B, D) Heatmap of the indicated differentially expressed genes from Lineage 1 - EryP (B) and Lineage - 3 MyeloP (D) along pseudotime split by lean (left) and obese (right). E) Line plot depicting the Log(expression+1) of the indicated differentially expressed genes for each cell along pseudotime for Lineage 3 MyeloP. F) Bar plot of HLA-DR MFI for the indicated cell type from the spectral flow cytometry panel.

We identified 461 DEGs between the two groups in MyeloP cells that mapped to GO terms associated with cell activation, cytokine stimulus, transcription factor binding, and phagocytosis (Fig.3C, SuppTable2). Obesity was associated with robust upregulation of multiple proinflammatory genes, including alarmins (*S100A4*, *S100A10*, *S100A6*), interferon stimulated genes (*IFI16*, *ISG15*), and other inflammatory mediators (*TNFSF13B, MIF*) (Fig.3D). Antimicrobial genes *LYZ* and *ELANE* were also upregulated (Fig.3D). Conversely, several transcription factors were downregulated including *SOX4,* a developmental transcription factor that regulates stemness and progenitor cell differentiation(64), and *ATF3*, a stress induced antimicrobial gene (62) (Fig.3D). Interestingly, despite the upregulation of alarmins we observed a decrease in the expression of two factors of the pro-inflammatory AP-1 transcription factor complex, *FOS* and *JUN*(65) (Fig.3D). Additionally, we observed upregulation of *IRF8,* a transcription factor essential for the differentiation and function of myeloid cells as well as the production of interferon stimulated genes(66) (Fig.3D). Furthermore, we observed altered expression of adhesion molecules, with downregulation of *SELL* and upregulation of *ITGB2* (Fig.3D). Interestingly, HLA utilization shifted in the obese group, with increased expression of MHC class I (*MAMU-A*) and reduced expression of MHC class II (*MAMU-DRB1*) (Fig.3E). This shift in MHC class expression suggests functional consequences on terminally differentiated monocytes, therefore we confirmed this difference at the protein level by plotting the MFI of HLA-DR on terminally differentiated monocytes of the FBM from our spectral flow cytometry data. We confirmed significantly decreased protein levels of MHC class II HLA-DR on both classical and non-classical monocytes (Fig.3F). These data demonstrate that maternal obesity alters fetal HSPC fate by reshaping differentiation trajectories and transcriptional programs, leading to altered maturation and inflammatory skewing of the myeloid lineage.

### Maternal Obesity Impairs HSC Communication and Signaling Networks

To investigate how maternal obesity alters hematopoietic cellular communication within the FBM, we applied CellChat analysis to the scRNA-seq data. Maternal obesity was associated with a reduction in the total number and overall strength of inferred cell-cell interactions (Fig.4A). At the cluster-to-cluster level, the obese group exhibited fewer and weaker interactions compared with lean controls (Fig.4B). We next quantified the relative information flow, an aggregate measure of signal number and strength, for each pathway in each group (Fig.4C). We identified several signaling pathways predicted to be stronger in the lean group (THBS, ANNEXIN, CD39, PDGF, PTPRM, and BTL) and the obese group (ESAM, MPZ and CD45) (Fig.4C). We next examined the differential use of signaling pathways across individual hematopoietic populations between the two groups (Fig.4D). With maternal obesity, both ST-HSC and LT-HSC clusters exhibited increased signaling strength of ESAM and FN1 pathways with ST-HSC showing increased signaling with CD45, whereas MPP subsets demonstrated increased activation of MIF and SELL pathways (Fig.4D). The MyeloP population showed the most pronounced changes, with strong upregulation of RESISTIN, CSF, JAM, and ICAM signaling pathways in the obese group (Fig.4D).

**Figure 4:**
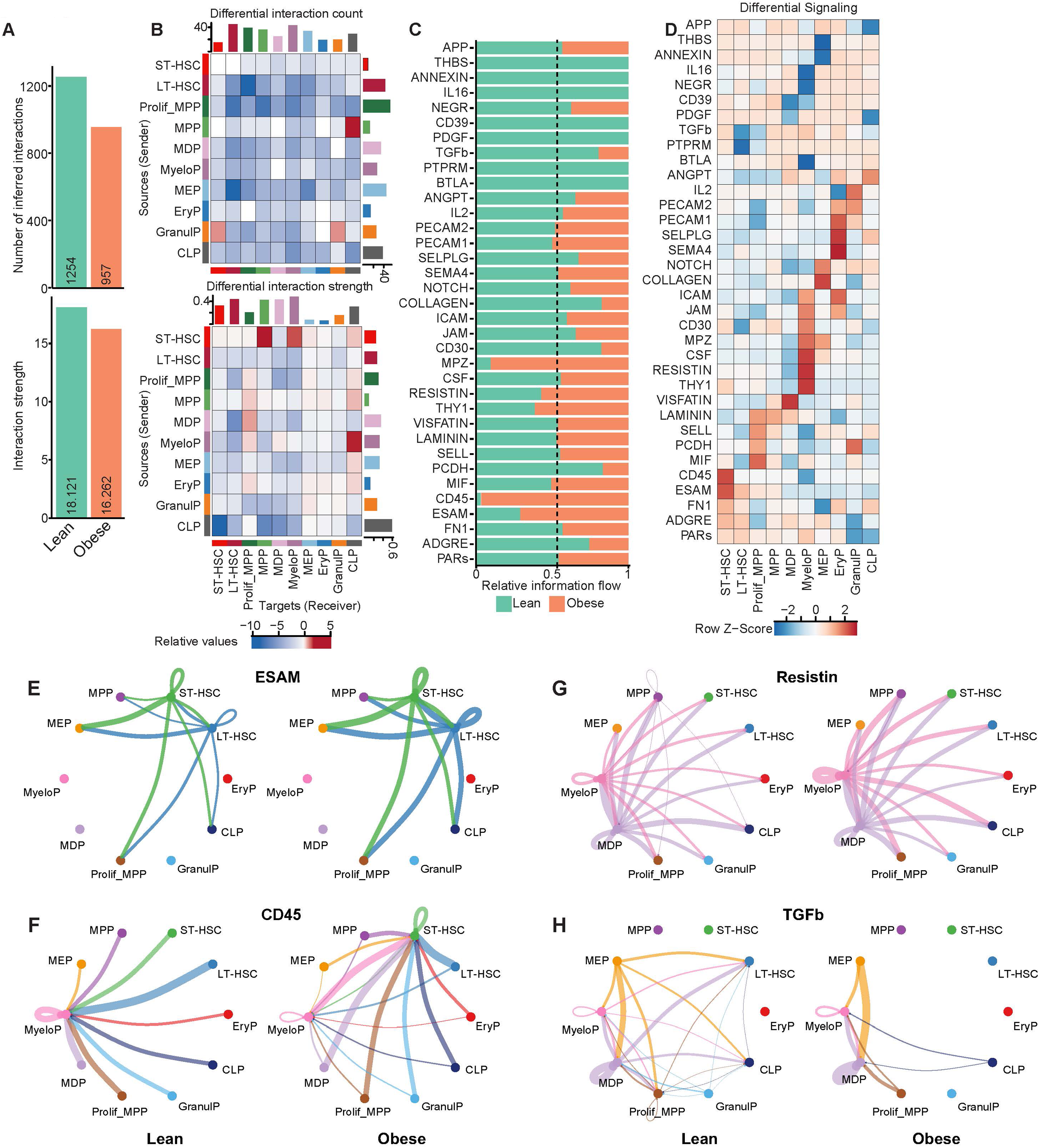
Altered signaling in FBM HSPCs indicates more rapid myelopoiesis in response to maternal obesity. A) Bar plot showing the inferred interactions (top) and strength (bottom). B) Heatmap showing the relative count (top) and interaction strength (bottom) between clusters. Blue indicates low interaction in the obese group compared to the lean group. Bars along the top or right side indicate the cumulative interaction of all signaling for the corresponding column or row, respectively. C) Bar plot of relative information flow (aggregate probability of communication) for each signaling pathway in FBM from lean (green) or obese (orange) groups. D) Heatmap showing the differential relative cluster-to-pathway interaction. Red indicates higher interaction potential in the obese group compared to the lean group. E-H) Circle plots showing the cluster-to-cluster interactions for the indicated pathways. The lean group is shown on the left and the obese group on the right. Line color indicates source of the interaction, and line thickness indicates the relative interaction count.

Pathway-level analysis of inter-cluster communication revealed several notable differences between the groups. Both HSC subsets exhibited pronounced upregulation of ESAM signaling to multiple clusters with maternal obesity, indicating dysregulation of a pathway essential for homeostatic hematopoiesis (Fig.4E) (67). We also report a complete shift in the target of CD45 signaling, with MyeloP cells being the primary recipient in the lean group shifting to the ST-HSC cluster with maternal obesity (Fig.4F). CD45 signaling in the bone marrow is essential for progenitor mobilization and egress(68); this shift suggests an increased demand for stem cell mobilization or regeneration in the FBM with maternal obesity. In addition, RESISTIN signaling, a proinflammatory pathway linked to enhanced myelopoiesis, was strongly upregulated in the MyeloP cluster in the obese group (Fig.4G) (69). Finally, maternal obesity resulted in a complete loss of TGFβ signaling received by the LT-HSC population and an increase in signaling to MyeloP and MDP cells (Fig.4H). Overall, maternal obesity disrupts FBM hematopoietic communication by weakening overall interaction strength, selectively amplifying pro-inflammatory and myelopoiesis-associated signaling pathways, and diminishing regulatory cues to HSC populations.

### Maternal Obesity Alters Monocyte Differentiation from HSPC and Their Function

Following the observation of pronounced changes in gene expression and cell-cell signaling within the MyeloP lineage, along with altered frequencies of terminally differentiated monocytes in the FBM, we assessed the capacity of sorted CD34+ HSPCs to differentiate into myeloid cells in vitro using the StemSpan myeloid supplement, which specifically promotes myeloid cell differentiation. Our findings indicate an accelerated differentiation of fetal CD34+ cells in the obese group, as evidenced by a significant increase in CD34-cells compared to the lean group (Fig.5A). Moreover, the frequency of the monocytes was increased after in vitro differentiation in the obese group (Fig.5B). Additionally, the mean fluorescent intensity (MFI) values of CCR2 and CD115 on monocytes was reduced (Fig.5C). Together this suggests that HSPCs in FBM exposed to maternal obesity are primed for rapid myeloid differentiation with potential impaired migratory and colony-stimulating capacity.

**Figure 5:**
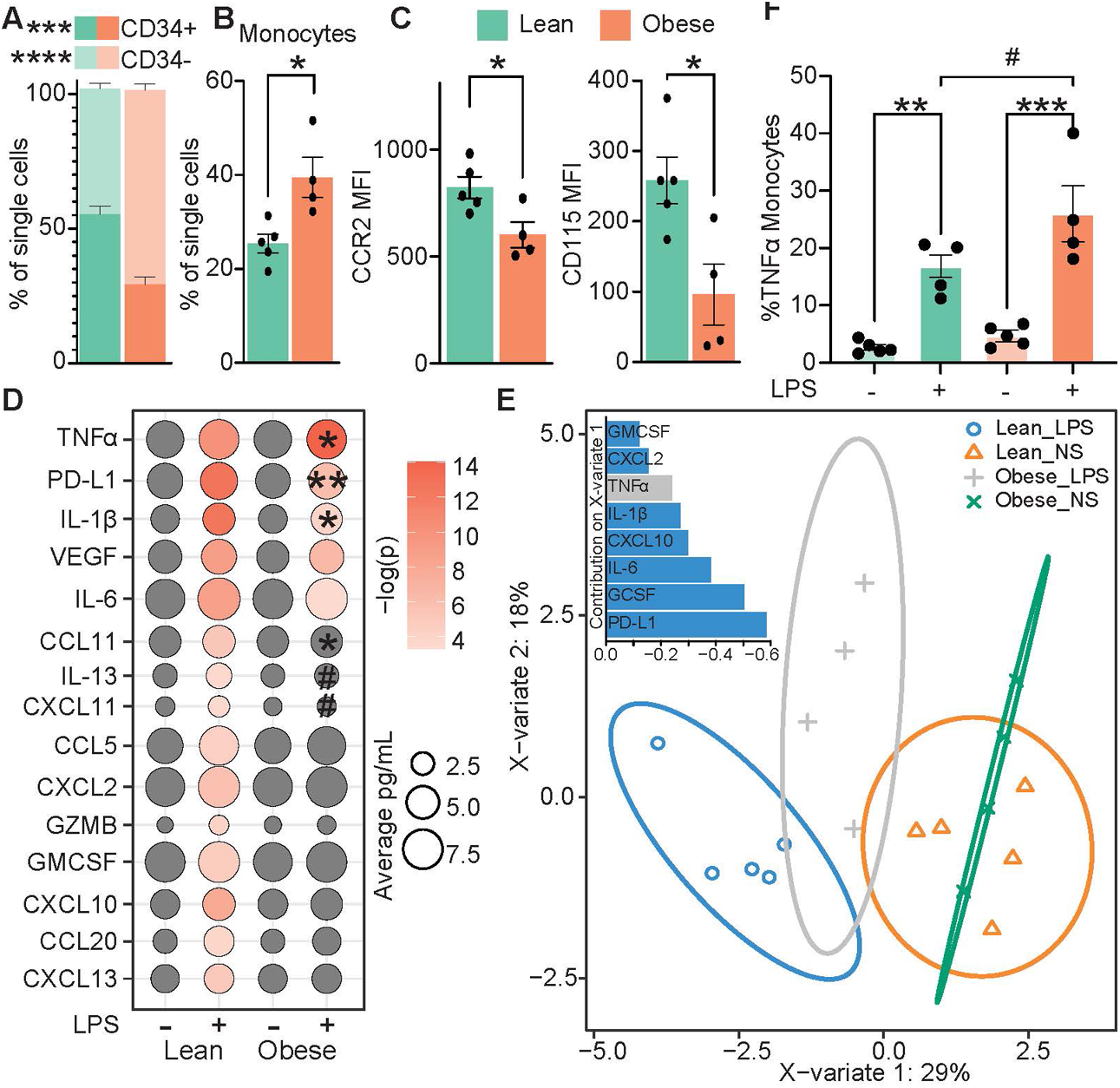
Maternal obesity induces functional alterations in in vitro differentiated HSPC-derived monocytes. A) Stacked bar graph showing CD34- and CD34+ cell frequencies following liquid differentiation assay. B) Bar graphs showing the frequency of CD14+HLA-DR+ Monocytes within the single-cell population following liquid differentiation assay. C) MFI of CCR2 (left) and CD115 (right) on Monocytes. D) Bubbleplot of analyte production after LPS stimulation. Size of the bubble indicates analyte concentration (pg/mL), and color represents the -log2(P-value) for the comparison between stimulated and non-stimulated conditions within each group (one-way ANOVA). E) sPLS-DA and inlayed bar plot of factors defining the variate 1 axis. F) Bar graph showing the percent of total monocytes expressing TNFα after stimulation with LPS.

We next interrogated the ability of differentiated cells to respond to LPS stimulation. Using a Luminex assay, we observed a robust increase in the production of 15 cytokines, chemokines and growth factors with stimulation in the lean group (Fig.5D). With stimulation, maternal obesity exposure was associated with increased levels of only five analytes: TNFα, PDL1, IL1β, VEGF, and IL6 (Fig.5D). However, the magnitude of this increase differed across analytes. Although PD-L1 and IL-1β were upregulated following stimulation in the obese group, their levels remained significantly lower than in the stimulated lean group. In contrast, TNFα showed the opposite pattern, with significantly higher concentrations after stimulation in the obese group compared to the lean group (Fig.5D). We next used the sparse partial least squares discriminant analysis (sPLS-DA), a supervised method that identifies the most predictive variables contributing to group separation, to determine which factors were driving the observed differences (70). The sPLS-DA sample plot demonstrated robust clustering of the sample groups, with variate 1 separating samples according to stimulation condition (Fig.5E). In the lean LPS group, stimulation-associated separation was primarily driven by GM-CSF, CXCL2, IL-1β, CXCL10, IL-6, G-CSF, and PD-L1, whereas in the maternal obesity group, TNFα emerged as the main driver of stimulation-induced variation. (Fig.5E). As the Luminex measures the analyte production from the total population of differentiated cells, we next validated the TNFα response using intracellular cytokine staining, gating specifically on monocytes (CD14+HLA-DR+). Consistent with the Luminex results, both groups responded to stimulation; however, the magnitude of this response was significantly higher with exposure to maternal obesity. (Fig.5F).

### Maternal Obesity Drives Expansion of Intermediate Progenitor Populations and Alters Lineage Progression

Given the observed changes in HSC frequency and total monocyte output following in vitro differentiation, along with the limited functional capacity of the differentiated cells, we next examined the impact of maternal obesity on HSC differentiation at single-cell resolution using scRNA-seq after StemSpan differentiation. Data integration from both lean and obese groups identified nine distinct cell clusters (Fig.6A). A small population of HSCs was defined by expression of *CD34*, *HOPX* and *CD38* (Fig.6A,B and SuppTable3). Two MPP populations were distinguished by expression of *CLEC12A, CD14*, *FCGR3*, and *CYBB*, with one cluster showing higher proliferative activity marked by *MKI67* expression (Fig.6A,B). GMP were defined by expression of *CSF1R, CCR2, and PTPRC* (Fig.6A,B), while a terminally differentiated myeloid population was characterized by high expression of *FCGR3*, *S100A11*, and *SIGLEC1* (Fig.6A,C). The neutrophil population was distinguished by absence of *CD14* and high expression of *FGR* and *S100A8/9* (Fig.6A,B). Interestingly, despite using the StemSpan Myeloid Supplements we detected the presence of two lymphoid progenitors. The CLP1 cluster expressed *CD74*, *MS4A1*, and *CD1C,* likely representing B cell progenitors, while the CLP2 cluster expressed *NKG2D*, *BCL11A*, *DNTT*, *IL7R*, and *CCR7*, likely representing T cells progenitors (Fig.6A,B). Finally, a megakaryocyte progenitor (MKP) cluster was defined by *HBA*, *PLEK*, and *GPX4* expression (Fig.6A,B).

**Figure 6:**
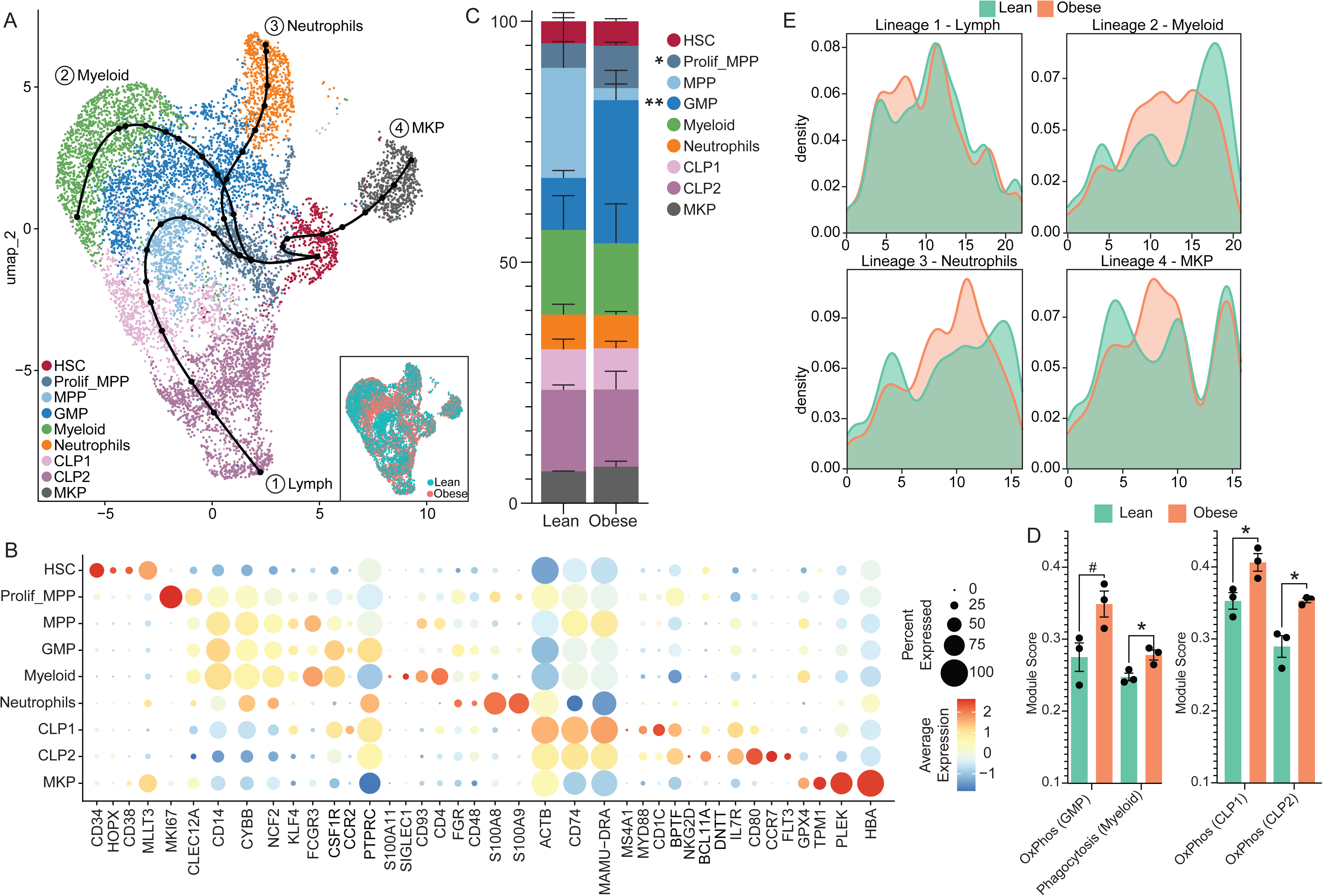
FBM HSPCs from obese dams remain in an intermediate pluripotent state. A) UMAP of 10,053 sorted fetal CD34+ cells (5,775 lean and 4,278 obese) from the lean (n=3) or obese (n=3) groups, following differentiation in StemSpan media for 14 days. UMAP is overlaid with four lineage trajectories. Inlayed UMAP colored by cell origin of fetuses from lean or obese dams, showing proper dataset integration. Colors indicate the nine distinct clusters. B) Bubble plot showing marker genes used to identify the nine clusters. Bubble size indicates the percentage of cells in each cluster per indicated marker. Bubble color represents the average marker expression across cells within the cluster. C) Stacked bar plot showing the relative cluster frequencies. D) Bar plot showing the indicated module score for the indicated clusters. E) Line plot depicting cell densities along pseudotime of the indicated trajectories.

Cluster frequency analysis revealed a significant expansion of Prolif_MPP and GMP populations in the obese group (Fig.6C), with a trend toward reduced MPP population (Fig.6C). Module scoring further revealed a trending increase in oxidative phosphorylation (OxPhos) in the GMP population and a significant upregulation of phagocytosis module scores in myeloid cells (Fig.6D). For both CLP groups, we saw significant OxPhos upregulation in the obese group compared to lean controls (Fig.6D). No other differences in module scores associated with hypoxic signatures, glycolysis or OxPhos were observed in any other cluster. Trajectory analysis, rooted in the HSC cluster, identified four unique lineages (Fig.6A). Lineage 1 progressed through Prolif_MPP, into MPP, then through both CLP populations. (Fig.6A and Supp.Fig3A). Lineages 2 and 3 branched from proliferative MPP into GMP, terminating in myeloid (lineage 2) or neutrophil (lineage 3) clusters (Fig.6A and Supp.Fig3B-C). Lineage 4 transitioned directly from HSC to the MKP cluster (Fig.6A and Supp.Fig3D). Across all four lineages, pseudotime analysis revealed significantly altered differentiation paths with obesity, characterized by aberrant accumulation of cells at mid-differentiation states (Fig.6E).

Maternal obesity also altered gene expression dynamics along the lymphoid and myeloid lineages, but not neutrophil and MKP lineages (SuppTable4). The 642 DEGs along the lymphoid lineage enriched to GO terms associated with proinflammatory and humoral responses, cell migration, response to hypoxia, and cell killing (Fig.7A, SuppTable4). Notably, expression levels of *CCL17*, *CCL22*, *IL23A*, and *TNFRSF18* genes were increased, while expression of the inhibitory receptor *FCGR2B* was decreased in the obese group (Fig.7B). Along the myeloid trajectory (Lineage 2), we identified 183 DEGs that enriched to GO terms associated with response to bacterium, cell adhesion or chemoattractant activity, phagocytosis, complement activation, and pattern recognition receptor pathways (Fig.7C, SuppTable4). Proinflammatory genes, including *IFI30*, *CXCL16,* and *IL1B,* were consistently upregulated with maternal obesity (Fig.7D). Furthermore, there is increased expression of *CD80,* an activation marker that aids in antigen presenting cell signaling to T cells (71), and *CD36,* a scavenger receptor required for functional phagocytosis(72), whereas *FCGR2B* expression was decreased (Fig.7D). Additionally, *CCL2* was constitutively upregulated, while *CCL7* expression declined rapidly in myeloid cells from the obese group (Fig.7E). Collectively, our findings indicate that maternal obesity reprograms HSPC differentiation toward proliferative, myeloid-biased lineages, with a concomitant accumulation of intermediate progenitors and upregulation of proinflammatory transcriptional programs.

**Figure 7:**
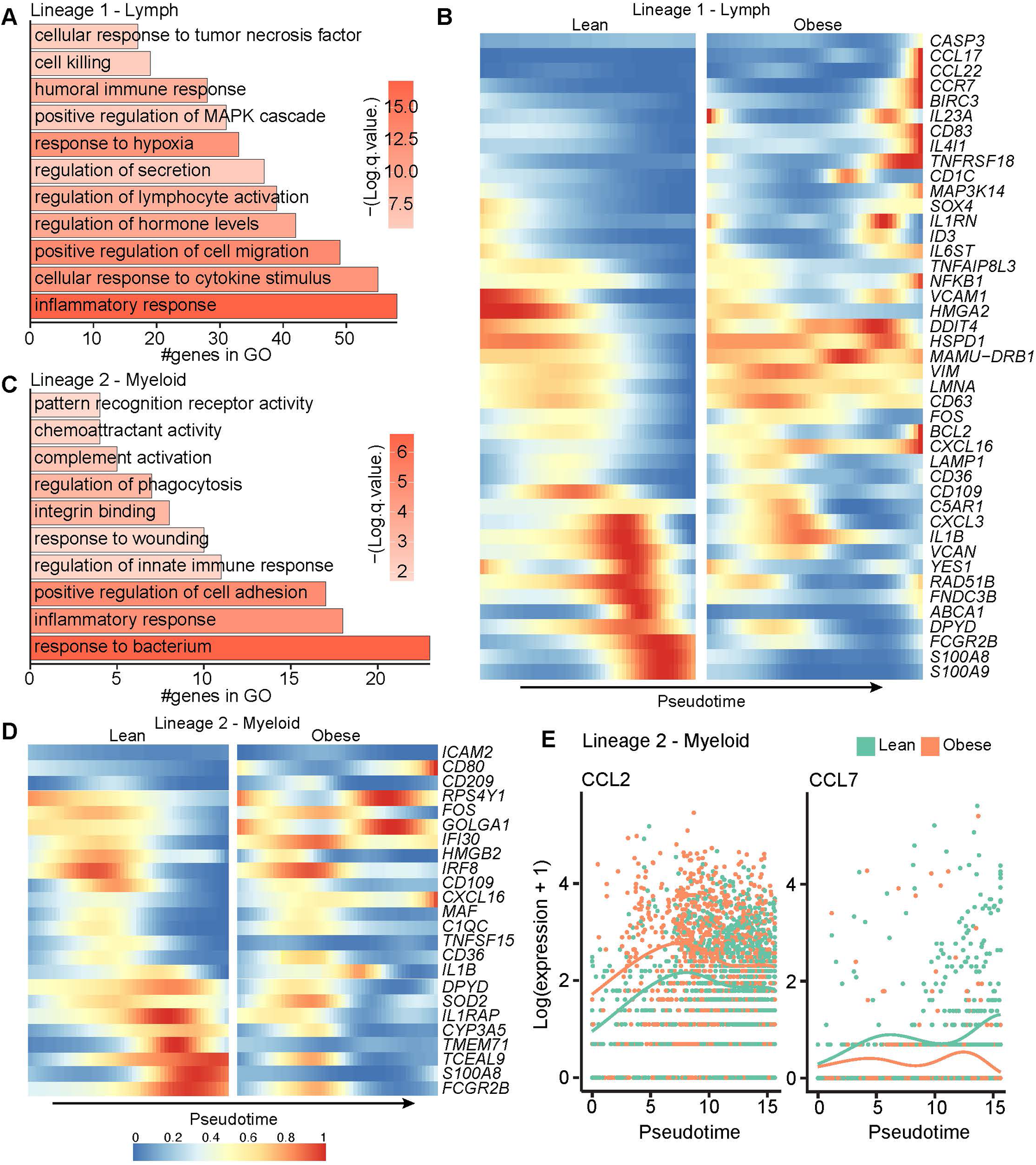
Trajectory analysis of differentiated FBM HSPCs demonstrates altered transcriptional profile of myeloid and lymphoid cells. A, C) Bar plot of gene ontology terms for genes differentially expressed along pseudotime in Lineage 1 - Lymph (A) and Lineage 2 - Myeloid (C). Bar length represents the number of genes mapping to each term and bar color indicates the -log2(Q-value). B, D) Heatmap of the indicated differentially expressed genes from Lineage 1 - Lymph (B) and Lineage 2 - Myeloid (D) along pseudotime split by lean (left) and obese (right). E) Line plot depicting the Log(expression+1) of the indicated differentially expressed genes for each cell along pseudotime for Lineage 2 Myeloid.

## Discussion

Maternal obesity has been linked to altered immune system development and function in offspring (12–17, 20–23). Recent findings indicate that maternal obesity alters HSPC fate decision and lineage commitment, as well as the cellular composition of the bone marrow niche (27–31). Although prior work has begun to explore the impact of maternal obesity on the fetal immune development (15–17, 20–23), clinical studies are inherently limited in mechanistic insight due to restrictions on postnatal sample collection and ethical barriers to obtaining fetal tissues (20–23). Some of these limitations can be addressed using animal models; however, many rely on obesogenic diets that may not fully recapitulate human obesity, which arises from a combination of dietary patterns, lifestyle factors, and genetic predisposition (34, 35). To address this knowledge gap, we examined the impact of maternal obesity on FBM HSPC phenotype using lean or obese rhesus macaques selected based on body composition, thereby representing individuals with potential genetic predisposition to obesity.

HSC in the bone marrow balance self-renewal with differentiation into all mature blood lineages(24–26). HSC are comprised of LT-HSC, which are more quiescent and possess extensive self-renewal capacity to sustain lifelong multilineage hematopoiesis, and ST-HSC which have limited self-renewal capacity and differentiate rapidly to produce mature blood cells(47, 48). We observed no difference in the frequency of LT-HSC; however, the abundance of ST-HSC was increased with maternal obesity. This finding suggests that maternal obesity may limit the long-term stemness of HSC, transcriptionally predisposing them toward differentiation. This idea is in line with our flow cytometry data that maternal obesity leads to reduced frequencies of primitive HSC and our previous study in rhesus macaques showing that maternal obesogenic WSD leads to reduced functional capacity of FBM HSPCs in vivo (73). Furthermore, we identified dysregulation of key signaling pathways with major implications for the balance between self-renewal and differentiation of HSC. We observed heightened signaling along the endothelial cell-selective adhesion molecule (ESAM) pathway in both HSC populations. In murine studies, the ESAM pathway has been identified as a key regulator of hematopoiesis, with ESAM expression elevated on actively dividing HSC and multipotent myeloid-erythroid progenitors (67). We also observed a loss of TGFβ signaling in the LT-HSC population with maternal obesity. This growth factor has been shown to be a key regulator of HSC quiescence and a potent inhibitor of HSC differentiation (74). Loss of this signaling pathway to the LT-HSC population further indicates an inability to maintain self-renewal. In contrast, we observed increased signaling of TGFβ to myeloid progenitors (MDP and MyeloP) and it has been shown that higher levels of this growth factor in early progenitors have been linked to a shift in hematopoiesis from lymphopoiesis toward myelopoiesis (75). The observed alteration of ESAM and TGFβ signaling skew hematopoiesis toward the myeloid lineage, suggesting that maternal obesity may transcriptionally skew HSC towards myeloid differentiation. These findings are consistent with prior studies where *ESAM* and *TGFB1* gene expression was significantly decreased in the FBM of WSD exposed fetal rhesus macaques and in mouse models of HFD, where fetal HSC are skewed toward myelopoiesis (27–30, 76).

We observed an increased frequency of GMP populations by flow cytometry and after in vitro differentiation by scRNA-seq, with an increased total number of CD14+HLA-DR+ monocytes following *in vitro* differentiation. Additionally, we observed a significant decrease in CCR2 and CD115 by flow cytometry, as well reduced expression and signaling of genes important for migration and adhesion, including *SELL* on monocytes (Fig.3D and 4D). While CD115 is essential for the survival, proliferation, and differentiation of myeloid cells, CCR2 plays a critical role in the egress of monocytes from the bone marrow into the periphery (77). Studies have shown that CCR2-deficient mice exhibit increased monocyte retention in the bone marrow and decreased monocyte levels in the circulation (78). We have previously shown that umbilical cord blood monocytes are decreased in human neonates born to mothers with obesity (22). Monocyte egress from the bone marrow is initiated by CCR2 activation triggered by CCL2 released by stromal cells or CCL7 produced primarily by monocytes and macrophages (77, 78). In our trajectory analysis of differentiated cells, we observed increased CCL2 expression across the entire pseudotime of myeloid differentiation with maternal obesity. However, we also observed a decrease in CCL7 expression along pseudotime with maternal obesity. Taken together, these findings indicate that maternal obesity may impair monocytes’ egress, potentially via disruption of the CCR2-CCL7 axis.

Additionally, we observed an upregulation of RESISTIN signaling within FBM myeloid progenitor cells exposed to maternal obesity. RESISTIN is primarily produced by adipocytes and monocytes, and upon interacting with TLR4, initiates proinflammatory signaling cascades involving NF-κB and MAPK that have been associated with enhanced myelopoiesis (69). Indeed, we observed increased expression of *MAP3K1* and various proinflammatory genes. Additionally, we observed decreased protein (MFI) and gene expression levels of MHC class II in monocyte progenitors with maternal obesity. Further, we observed the inability of *in vitro* differentiated cells to mount a response to ligand stimulation. Loss of MHC class II expression on monocytes leads to defective antigen presentation, resulting in impaired activation of CD4+ T cells and a shift toward an immune-tolerant or dysfunctional state, thereby compromising effective immune defense and pathogen clearance (79).

In addition to the observed heightened inflammatory transcriptional state of terminally differentiated myeloid cells, indicated by upregulated activation markers and cytokines, we also observed increased scores for glycolysis and phagocytosis modules. Glycolysis is a metabolic state associated with cellular inflammation and immune activation (80). This contrasts with the reduced glycolysis observed in monocytes from human UCB with maternal obesity (22), suggesting distinct metabolic mechanisms in monocytes before and after egress from the FBM. Additionally, the observed increase in phagocytic potential aligns with findings in monocytes form human UCB with maternal obesity (22), and heightened phagocytic activity has been associated with more regulatory monocyte/macrophage phenotypes (81).

Our findings reveal significant alterations in the transcriptional dynamics of lymphoid progenitors in response to maternal obesity, indicating a reprogramming of lymphoid differentiation kinetics and functionality. Temporal shifts in the expression of genes critical for autophagy and intracellular trafficking (*AP4E1, NFKB1, LAMP1)* indicate that maternal obesity may disrupt intracellular communication, potentially altering immune competency later in life. Autophagy has been shown to be critical for T cell proliferation and maintenance of downstream signaling after TCR engagement(82, 83). Indeed, we have previously shown that pregravid obesity predisposes CD4+ T cells in human UCB toward a memory phenotype with limited functional capacity following ex vivo stimulation(20). We also observed transcriptional changes in lymphoid progenitors indicative of potential skewing of the lymphoid lineages toward more innate-like NKT cells. Specifically, downregulation of *EAF2*, an elongation factor regulating proliferation and apoptosis in B cells, which deficiency causes excess B cell death and hyperactivation(84, 85); *BCAT1*, a gene induced by BCR/TLR9 co-activation and is essential for B-cell growth and survival(86); and *UGCG,* which regulates glycosphingolipid biosynthesis and is necessary for the formation of cytotoxic granules and clonal expansion of cytotoxic T cells after infection (87, 88). These data are in line with our previous studies in macaques that reported reduced lymphopoiesis and B cell frequencies in FBM with maternal WSD (73). In contrast, expression of *NRP1*, a membrane receptor that has been shown to be a marker of NKT cells in the thymus (89, 90) and *ZBTB16,* a transcription factor essential for the differentiation and intrathymic expansion of early invariant NKT cells progenitors and influences their migration to the periphery(91, 92) were upregulated. Interestingly, it has been shown that maternal obesity is associated with increased frequencies of NKT cells in human UCB (21). Additionally, pDCs derived from the lymphoid lineage were decreased. These findings suggest that maternal obesity impacts HSPC function even in the absence of an obesogenic diet.

The FBM niche contains a non-hematopoietic (Lin-CD34-) compartment enriched in mesenchymal stromal/stem and progenitor cells (MSPCs), osteoblasts, osteoclasts, and adipocytes, which collectively help shape the microenvironment required for HSC survival, proliferation, and differentiation (26). While the individual frequency of these populations was not measured directly, we observed an expansion of the total Lin-population despite the decreased frequency of HSPC populations likely indicating an expansion of this non-hematopoietic compartment. This hypothesis aligns with previous reports of altered MSPC differentiation, resulting in more numerous and larger adipocytes in the FBM with maternal obesity (30, 31), and offers a possible mechanism underlying impaired HSPC function. Expansion of the non-hematopoietic fraction of FBM has been shown to limit HSPC populations by both restricting physical niche space and altering cellular signals that support stem cell function (93–95). Additionally, expansion of the CD34-fraction is consistent with a shift toward a proinflammatory FBM microenvironment. It has been shown in mice that obesity-induced inflammatory cues, particularly IL-1β, drive the expansion of the stromal cell populations(96, 97). Here, we report an upregulation of multiple proinflammatory genes (*S100A4*, *IFI16*, *ISG15*, *TNFSF13B* and *MIF)* in HSPCs with maternal obesity. This pro-inflammatory environment has been linked to increased HSC apoptosis and reduced self-renewal capacity (98, 99), which could underlie the decrease in HSC frequencies observed by flow cytometry and the apparent expansion of ST-HSCs, reflecting diminished long-term self-renewal potential.

The present study has some limitations. Our scRNA-seq analysis was restricted to CD34+ HSPC, which limited our ability to fully assess the abundance and cellular interactions of non-hematopoietic or terminally differentiated fractions of the FBM. Similarly, although high-parameter spectral flow cytometry allowed characterization of diverse HSPC and terminally differentiated populations, our panel lacked the markers necessary to further delineate the Lin⁻CD34⁻ compartment. Additionally, the study is limited to FBM. Nevertheless, in conclusion, our findings reveal that maternal obesity remodels the FBM microenvironment by promoting a proinflammatory transcriptional state, expanding non-hematopoietic populations, skewing HSPC differentiation toward myelopoiesis, and impairing immune cell responses to antigenic stimulation.

## Supporting information

SuppTable1

SuppTable2

SuppTable3

SuppTable4

## Funding

This study was funded by R01AI142841(I.M.) and P51 OD011092 for operation of the ONPRC.

## Data Availability

Data underlying the transcriptome analysis is available in the SRA under project number PRJNA1247568.

## Author contributions

O.V. and I.M. conceived the idea. B.M.D., H.H., S.B.W., and M.B.B. completed all experiments. B.M.D. performed the analysis. B.M.D., O.V., and I.M. interpreted the data and prepared the manuscript. All authors read and approved the final manuscript.

## Acknowledgments

We thank members of the veterinary/time mated breeding staff Lauren Drew Martin, and Travis Hodge at the Oregon National Primate Center at the Oregon for support with animal care and breeding. This work was supported by the UK Flow Cytometry & Immune Monitoring core facility. (RRIDSCR_026358).

## Declaration of interests

All authors declare no competing interests.

## Figure legends

**Supp Table 1: CD34+ Bone Marrow Marker genes**

**Supp Table 2: CD34+ Bone Marrow Trajectory DEGs**

**Supp Table 3: Liquid Differentiation Marker genes**

**Supp Table 4: Liquid Differentiation Trajectory DEGs**

## Abbreviations

BMI: body mass index
UCB: umbilical cord blood
FBM: fetal bone marrow
HFD: high fat diet
WSD: western style diet
GD: gestational day
HSC: hematopoietic stem cell
HSPC: hematopoietic stem and progenitor cell
MPP: multipotent progenitors
MEP: megakaryocyte and erythrocyte
MLP: multi lymphoid progenitors
CLP: common lymphoid progenitors
CMP: common myeloid progenitors
cMoP: common monocyte progenitors
GMP: granulocyte–monocyte progenitors
pDC: plasmacytoid dendritic cells
ST-HSC: short-term hematopoietic stem cell
LT-HSC: long-term hematopoietic stem cell
MDP: monocyte dendritic cell progenitor
MyeloP: myeloid progenitors
EryP: erythroid progenitors
GranulP: granulocyte progenitors

---

**Supp Figure 1:**
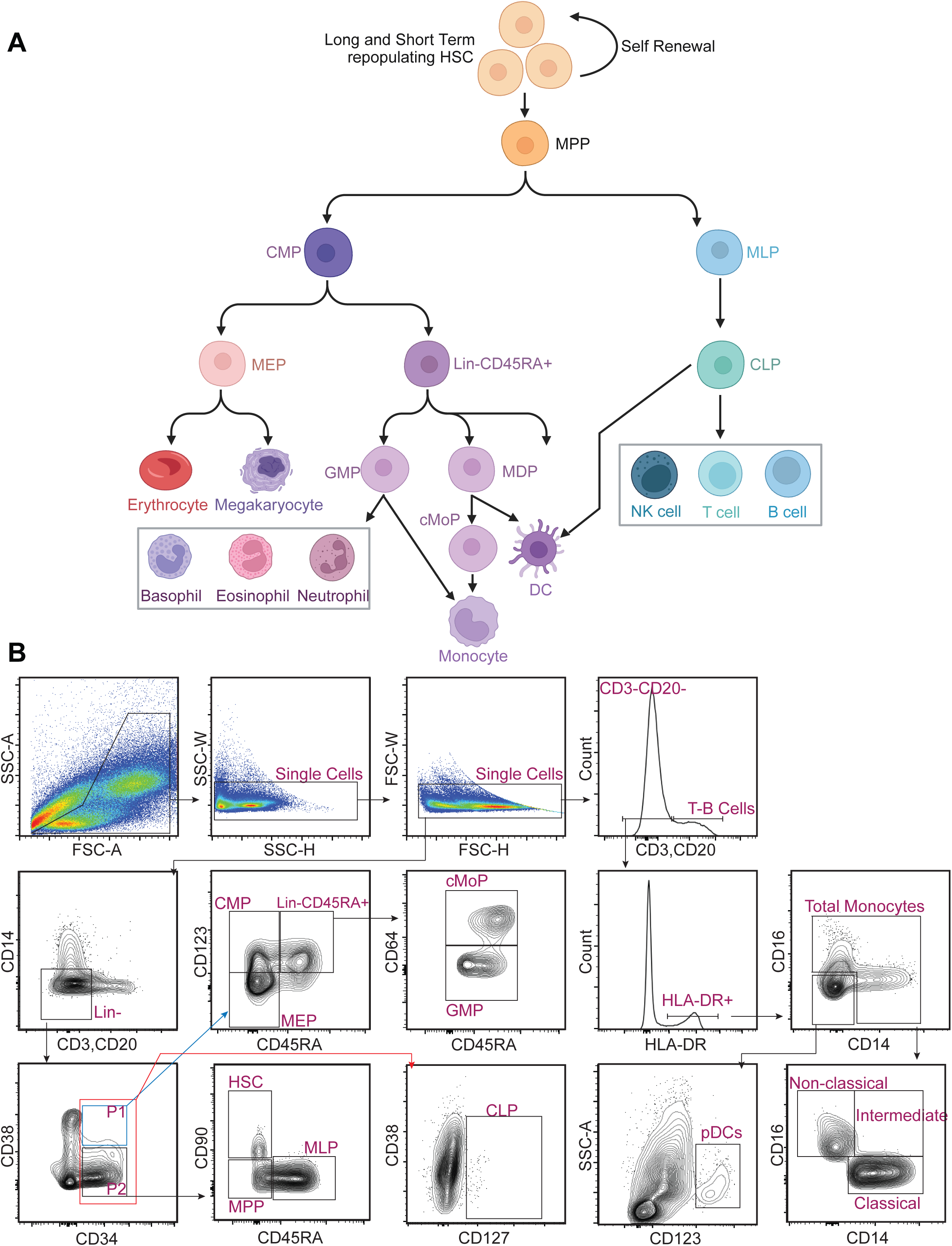
Spectral flow cytometry of FBM. A) Hematopoietic tree of cell differentiation. Image generated in BioRender. B) Gating of flow data using in Flow Jo for the analysis of various cell hematopoietic populations.

**Supp Figure 2:**
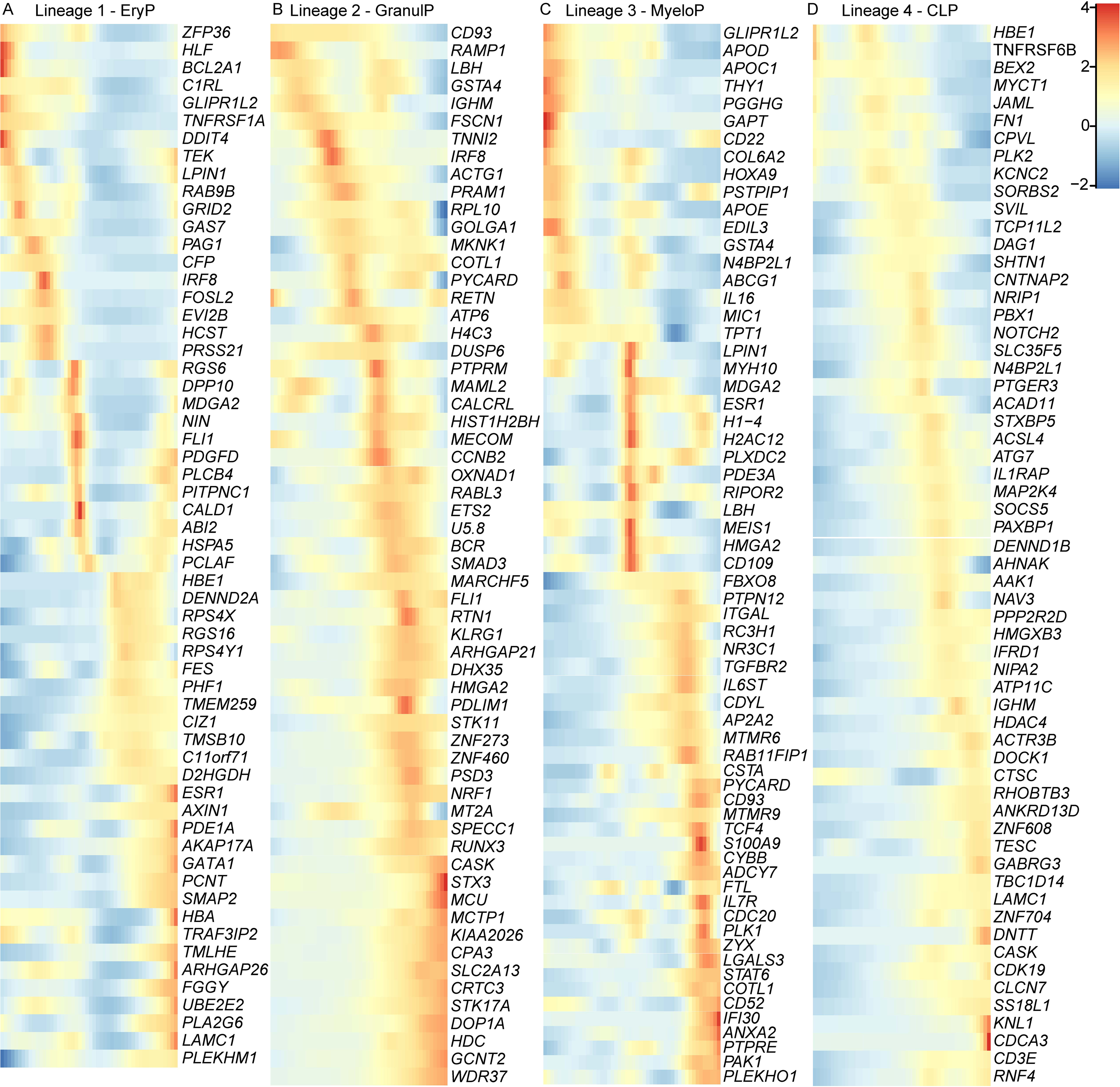
Defining trajectory lineages in CD34+ FBM cells. Heatmap of genes along pseudotime used by Slingshot to define the trajectories for A) Lineage 1 EryP, B) Lineage 2 GranulP, C) Lineage 3 MyeloP, and D) Lineage 4 Lymph.

**Supp Figure 3:**
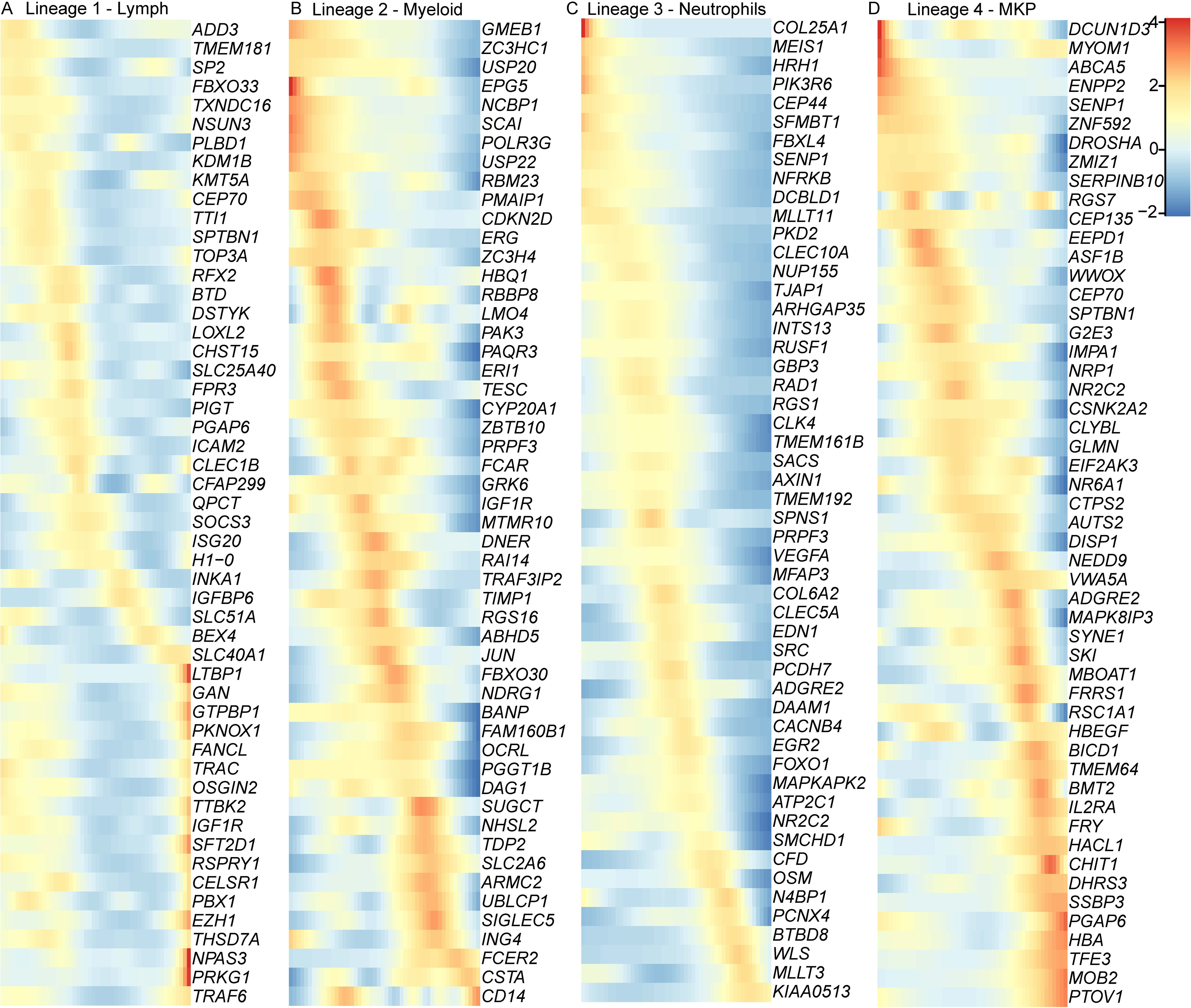
Defining trajectory lineages in CD34+ FBM cells after in vitro differentiation. Heatmap of genes along pseudotime used by Slingshot to define the trajectories for A) Lineage 1 Lymph, B) Lineage 2 Myeloid, C) Lineage 3 Neutrophils, and D) Lineage 4 MKP.

## Notes

### Competing Interest Statement

The authors have declared no competing interest.

